# Prediction of inter-chain distance maps of protein complexes with 2D attention-based deep neural networks

**DOI:** 10.1101/2022.06.19.496734

**Authors:** Zhiye Guo, Jian Liu, Jeffrey Skolnick, Jianlin Cheng

## Abstract

Residue-residue distance information is useful for predicting the tertiary structures of protein monomers or the quaternary structures of protein complexes. Many deep learning methods have been developed to predict intra-chain residue-residue distances of monomers accurately, but very few methods can accurately predict inter-chain residue-residue distances of protein complexes. We develop a new deep learning method CDPred (i.e., Complex Distance Prediction) based on the 2D attention-powered residual network architecture to address the gap. CDPred predicts the inter-chain distance maps of dimers (homodimers or heterodimers) from the features extracted from multiple sequence alignments (MSAs) and the intra-chain distance maps of predicted tertiary structures of monomers. Tested on two homodimer test datasets, CDPred achieves the precision of 61.56% and 43.26% for top L/5 inter-chain contact predictions (L: length of the monomer in homodimer), respectively, which is substantially higher than DeepHomo’s 37.40% and 23.08% and GLINTER’s 48.09% and 36.74%. And tested on the two heterodimer test datasets, the top L/5 inter-chain contact prediction precision (L: length of the shorter monomer in heterodimer) of CDPred is 47.59% and 22.87% respectively, which surpasses GLINTER’s 23.24% and 13.49%. Moreover, we demonstrate that the residue-residue co-evolutionary features calculated from multiple sequence alignments by a deep learning language model are more informative for the inter-chain contact prediction than the traditional statistical optimization approach of maximizing direct co-evolutionary signals, and large intra-chain distances in the intra-chain distance maps of monomers are more useful for the inter-chain distance prediction than small intra-chain distances.

## Introduction

Proteins are a key building block of life. The function of a protein is largely determined by its three-dimensional structure (Utsumi and Matsumura, 1997). Sometimes single-chain proteins (monomers) can perform certain functions, while the structures of most individual proteins interact to form multi-chain complex structures (multimers) to carry out their biological function (Spirin and Mirny, 2003). Therefore, modeling the three-dimensional structure of both monomers and protein complexes is crucial for studying protein function.

Deep learning has been applied to advance the prediction of the tertiary structures of monomers since 2012 (Eickholt and Cheng, 2012). Over a decade, many deep learning methods were developed to predict intra-chain residue-residue contact maps or distance maps of monomers (Adhikari, et al., 2018; Li, et al., 2019; Senior, et al., 2020; Wang, et al., 2017; Wu, et al., 2020), which were used by contact/distance-based modeling methods such as CONFOLD (Adhikari and Cheng, 2018) and Rosetta (Rohl, et al., 2004) to build their tertiary structures. Extensive studies (Adhikari and Cheng, 2018; Kandathil, et al., 2022; Xu and Wang, 2019; Yang, et al., 2020) have shown that if a sufficiently accurate intra-chain distance map is predicted, then the protein’s tertiary structure can be accurately constructed. Most recently, AlphaFold2 (Jumper, et al., 2021) uses an end-to-end deep learning method to predict both tertiary structures and residue-residue distances of monomers, achieved a very high average accuracy (∼90 Global Distance Test (GDT-TS) score (Zemla, 2003) in the 14th Critical Assessment of Techniques for Protein Structure Prediction (CASP14) in 2020. Recently, AlphaFold2 was extended to AlphaFold-multimer(Evans, et al., 2021) and AF2Complex (Gao, et al., 2022) to improve the prediction of quaternary structures of multimers.

Following the deep learning revolution in the prediction of intra-chain residue-residue distances and tertiary structures, recently some deep learning methods were developed to predict the inter-chain residue-residue contact map of homodimers and/or heterodimers, such as DeepHomo (Yan and Huang, 2021), DRcon (Roy, et al., 2021), and GLINTER (Xie and Xu, 2022) that predicts the contact map for both homodimers and heterodimers using as input a graph representation of protein monomer structure and the row attention maps generated from multiple sequence alignments (MSAs) by the MSA transformer (Rao, et al., 2021). The attention map calculated by the MSA transformer is a kind of residue-residue co-evolutionary feature extracted from MSAs. It has been automatically trained on millions of the MSAs to capture the co-evolutionary information across many diverse protein families during its unsupervised pretraining. Despite the significant progress, the accuracy of inter-chain contact prediction is still much lower than that of intra-chain contact/distance prediction, which calls for the development of more methods to tackle this problem.

In this work, we develop a new protein complex distance prediction method (CDPred) based on a deep learning architecture combining the strengths of the deep residual network (He, et al., 2016), a channel-wise attention mechanism, and a spatial-wise attention mechanism to predict the inter-chain distance maps of both homodimers and heterodimers. As in GLINTER, the attention map of the MSA generated by the MSA transformer is used as one input for CDPred. The predicted distance map for monomers in dimers is used as another input feature. Different from the existing deep learning methods, CDPred predicts inter-chain distances rather than binary inter-chain contacts (contact or no contact) that the current methods such as DeepHomo and GLINTER predict. We test the CDPred rigorously on two homodimer test datasets and two heterodimer test datasets. For these datasets, CDPred yields much higher accuracy than DeepHomo and GLINTER.

## 2. Results

### 2.1 Evaluation of inter-chain contact prediction for homodimers

We compare CDPred with DeepHomo and GLINTER on the HomoTest1 homodimer test dataset with the results shown in **Table 1**. The input tertiary structures for all three methods are predicted structures corresponding to the unbound monomer structures. The results of DeepHomo are obtained from its publication. Three versions of CDPred are tested. The first version (CDPred_BFD) uses the MSAs generated from the BFD database as input. The second version (CDPred_Uniclust) uses the MSAs generated from the Uniclust30 database as input. The third version (CDPred) uses the average of the distance maps predicted by CDPred_BFD and CDPred_Uniclust as the prediction. Because DeepHomo and GLINTER predict binary inter-chain contacts at an 8 Å threshold instead of distances, we convert the inter-chain distance predictions of CDPred, CDPred_BFD, and CDPred_Uniclust into binary contact predictions for comparison.

**Table 1.** The precision of top 5, top 10, top L/10, top L/5, and top L contact predictions on the HomoTest1 test dataset for DeepHomo, GLINTER, and three versions of CDPred. L: sequence length of a monomer in a homodimer. Bold numbers denote the highest precision.

| Predictor | top 5 | top 10 | top L/10 | top L/5 | top L/2 | top L |
| --- | --- | --- | --- | --- | --- | --- |
| DeepHomo | - | 52.50 | 43.20 | 37.40 | 28.20 | - |
| GLINTER | 54.81 | 54.07 | 50.54 | 48.09 | 41.90 | 34.91 |
| CDPred_BFD | 63.57 | 62.50 | 61.24 | 58.26 | 52.78 | 47.08 |
| CDPred_Uniclust | 65.71 | 61.79 | 60.53 | 58.18 | 54.14 | 47.91 |
| CDPred | **68.57** | **66.43** | **64.55** | **61.56** | **55.87** | **50.06** |

CDPred achieves the highest contact prediction precision across the board among all the methods. For instance, CDPred has a top L/5 contact prediction precision of 61.56%, which is 24.16% percentage points higher than DeepHomo and 13.47% percentage points higher than GLINTER. CDPred performs better than both CDPred_BFD and CDPred_Uniclust, indicating that averaging the distance predictions made from the two kinds of MSAs can improve the prediction accuracy.

We also compared the methods above on the HomoTest2 homodimer test dataset (**Table2**). Similarly, CDPred performs best in terms of all the evaluation metrics. Combining the predictions of CDPred from two kinds of MSAs improves the prediction accuracy.

**Table 2.** The precision of top 5, top 10, top L/10, top L/5, and top L contact predictions on the HomoTest2 test dataset for DeepHomo, GLINTER, and CDPred predictors. Bold numbers denote the highest precision.

| Predictor | top 5 | top10 | top L/10 | top L/5 | top L/2 | top L |
| --- | --- | --- | --- | --- | --- | --- |
| DeepHomo | - | 30.43 | 27.32 | 23.08 | - | - |
| GLINTER | - | 43.04 | 40.18 | 36.74 | - | - |
| CDPred_BFD | 43.48 | 41.74 | 42.24 | 40.01 | 35.92 | 33.80 |
| CDPred_Uniclust | 48.70 | 45.22 | 43.32 | 39.64 | 37.46 | 32.32 |
| CDPred | **53.04** | **50.00** | **49.23** | **43.26** | **40.08** | **35.98** |

### 2.2 Evaluation of inter-chain contact prediction for heterodimers

We compare CDPred and a state-of-the-art heterodimer contact predictor GLINTER on both HeteroTest1 and HeteroTest2 heterodimer test datasets (see results in **Table 3** and **Table 4**, respectively). The input tertiary structures of monomers used by both methods are predicted by AlphaFold2. We use two different orders of monomer A and monomer B (AB and BA) in each heterodimer to generate input features for CDPred to make predictions. The average of the outputs of the two orders is used as the final prediction.

**Table 3.**
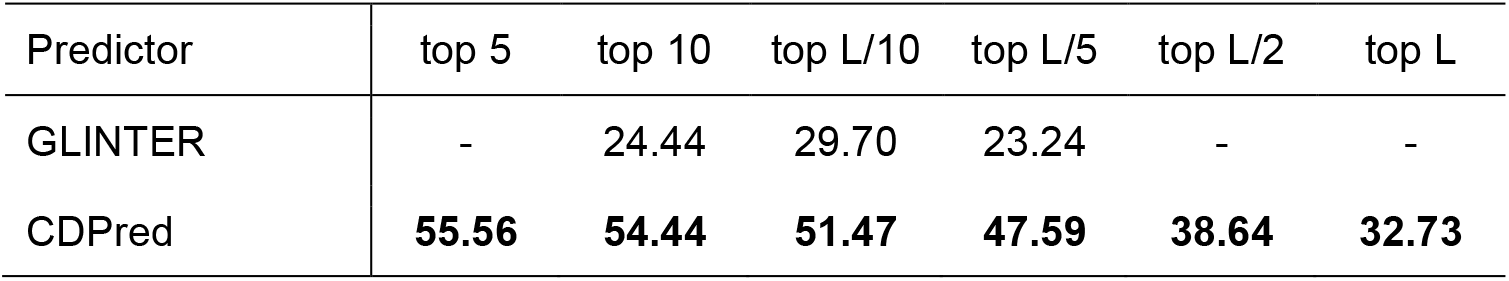
The precision of top 5, top 10, top L/10, top L/5, and top L contact predictions on the HeteroTest1 test dataset for the GLINTER and CDPred. L: the sequence length of the shorter monomer in a heterodimer. Bold numbers denote the highest precision.

**Table 4.** The precision of top 5, top 10, top L/10, top L/5, and top L contact predictions on the HeteroTest2 test dataset for the GLINTER and CDPred. L: the sequence length of the shorter monomer in a heterodimer. Bold numbers denote the highest precision.

| Predictor | top 5 | top 10 | top L/10 | top L/5 | top L/2 | top L |
| --- | --- | --- | --- | --- | --- | --- |
| GLINTER | 14.55 | 13.27 | 13.73 | 13.49 | 12.27 | 10.40 |
| CDPred | **23.27** | **23.82** | **23.93** | **22.87** | **20.17** | **17.51** |

On the HeteroTest1 dataset (**Table3)**, CDPred achieves much better performance than GLINTER in terms of all the metrics. For instance, the top L/5 contact prediction precision of CDPred 47.59%, is more than twice 23.24% that of GLINTER. On the HeteroTest2 dataset (**Table 4**), CDPred also substantially outperforms GLINTER.

Furthermore, we divide the top L/10 contact prediction precisions for the heterodimers in the more challenging HeteroTest2 dataset into four equal intervals and plot the number of heterodimers in each interval (**Figure 1**). The precision of the predictions in the four internals is bifurcated, mainly centered on a low precision interval [0%-25%] and a high precision interval [75%-100%]. 40 heterodimers have low contact prediction precision in the range of 0%-25%, indicating there is still a large room for improvement. One reason for the low precision is that most of the 40 heterodimers have shallow MSAs. The Pearson correlation coefficient between the logarithm of the number of effective sequences (Neff) of MSA and the top L/10 complex contact precision is 0.46, indicating a modest correlation between the two.

**Fig. 1.**
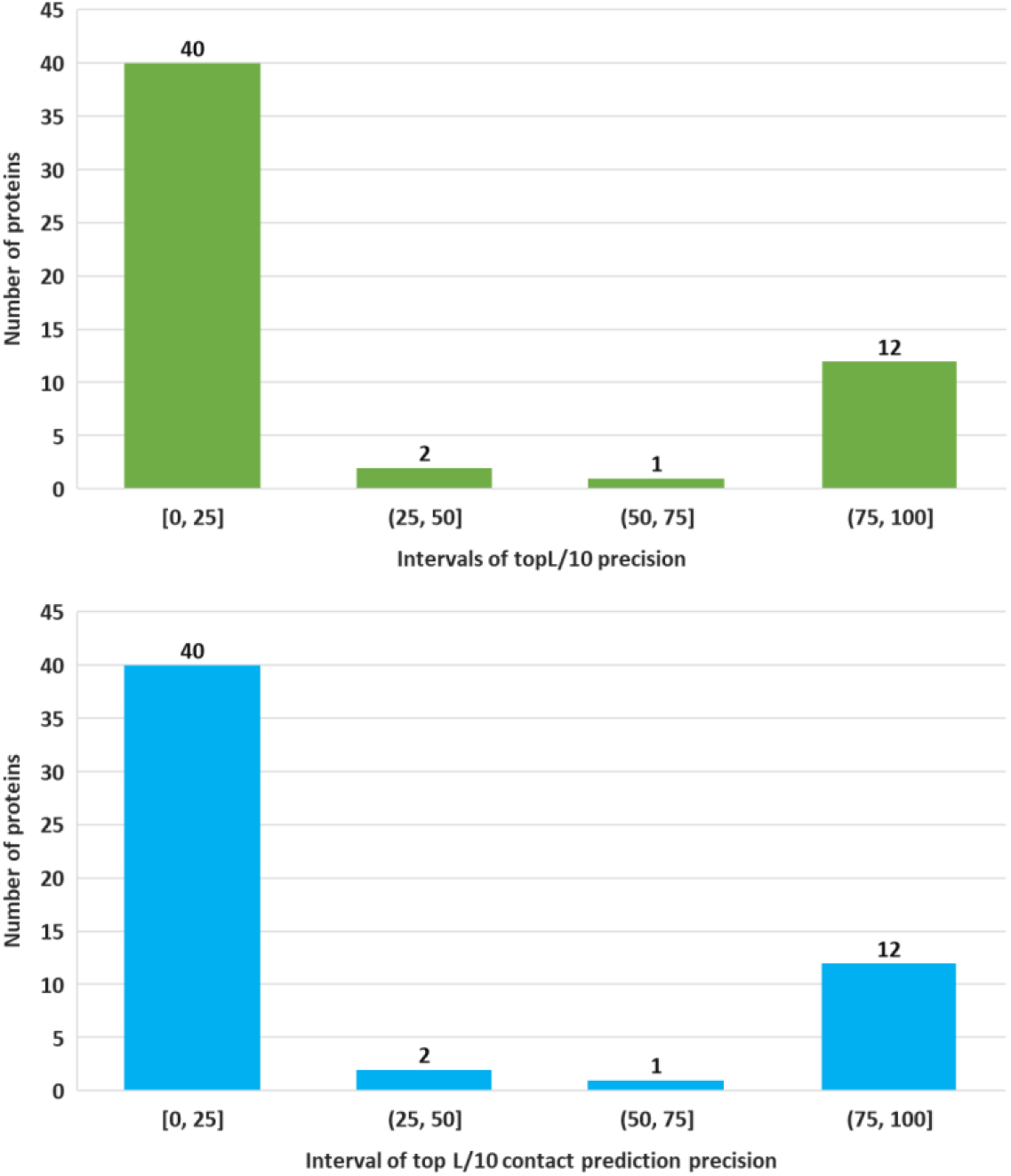
The histogram of the precision of the top L/10 contact predictions for the heterodimers in the HeteroTest2 dataset. The X-axis is the four precision intervals from 0% to 100%. The Y-axis is the number of heterodimers whose contact precision falls in each interval. Each interval has 40, 2, 1, and 12 heterodimers, respectively.

It is also observed that the inter-chain contact prediction accuracy for heterodimers is lower than for homodimers on average. One reason is that the MSA generation for a homodimer only needs to generate an MSA for a monomer in the homodimer, which is usually much deeper than the MSA generated for a heterodimer that requires the pairing of the related sequences in the MSAs of two different monomers in the heterodimer. Another reason is that homodimers tend to have a larger interaction interface than heterodimers on average, making the prediction easier.

### 2.3 Comparison of the co-evolutionary features generated by the statistical optimization method and deep learning method

To compare the performance of the co-evolutionary feature generated by the statistical optimization tool – CCMPred and the deep learning tool – MSA transformer, we trained two different models on the two different kinds of co-evolutionary features of the same training dataset using the same neural network architecture. One network (CDPred_PLM) is trained on the PLM co-evolutionary features generated by CCMPred. Another one (CDPred_ESM) is trained on the row attention map features generated by the MSA transformer. The precision of the top L/10 contact predictions of the two models on the four different test datasets are plotted in **Figure 2**. CDPred_ESM has better performance than CDPred_PLM on all the four test datasets, indicating that the co-evolutionary features extracted automatically by the deep learning method is more informative than by the statistical optimization method of maximizing direct co-evolutionary signals. However, combining the two kinds of co-evolutionary features yields even better results (see the results in **Tables 1, 2, 3**, and **4**). **Figure 3** plots the top L/10 precision of CDPred_ESM against the top L/10 precision of CDPred_PLM for the homodimers in the two homodimer test datasets and the heterodimers in the two heterodimers test datasets, respectively. For 42 out of 51 homodimers and 55 out of 64 heterodimers, CDPred_ESM has higher precision than CDPred_PLM. Both CDPred_ESM and CDPred_PLM can perform better on some targets, indicate the co-evolutionary features used by the two methods have some complementarity.

**Fig. 2.**
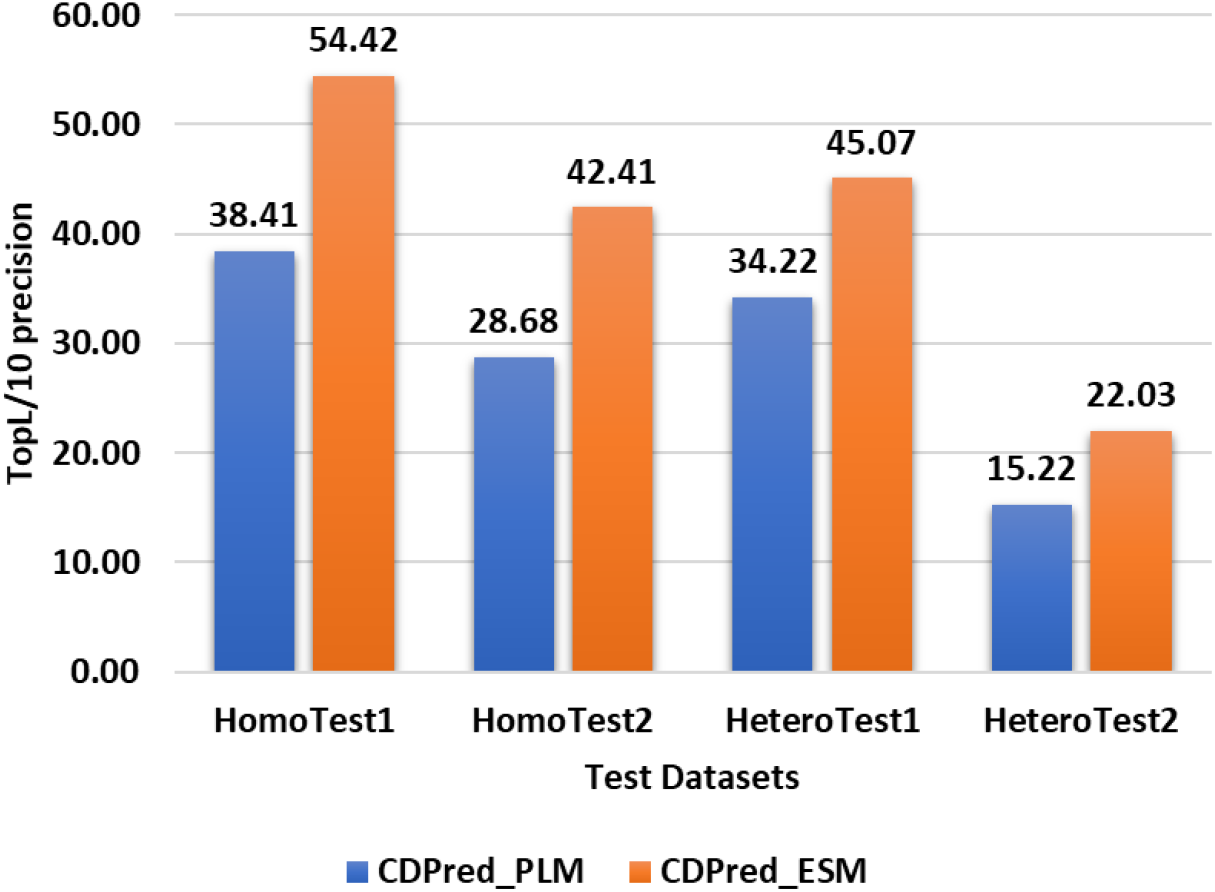
Comparison between CDPred_PLM and CDPred_ESM on four different test datasets. The y-axis is the top L/10 contact prediction precision, and the x-axis is the four different test datasets.

**Fig. 3.**
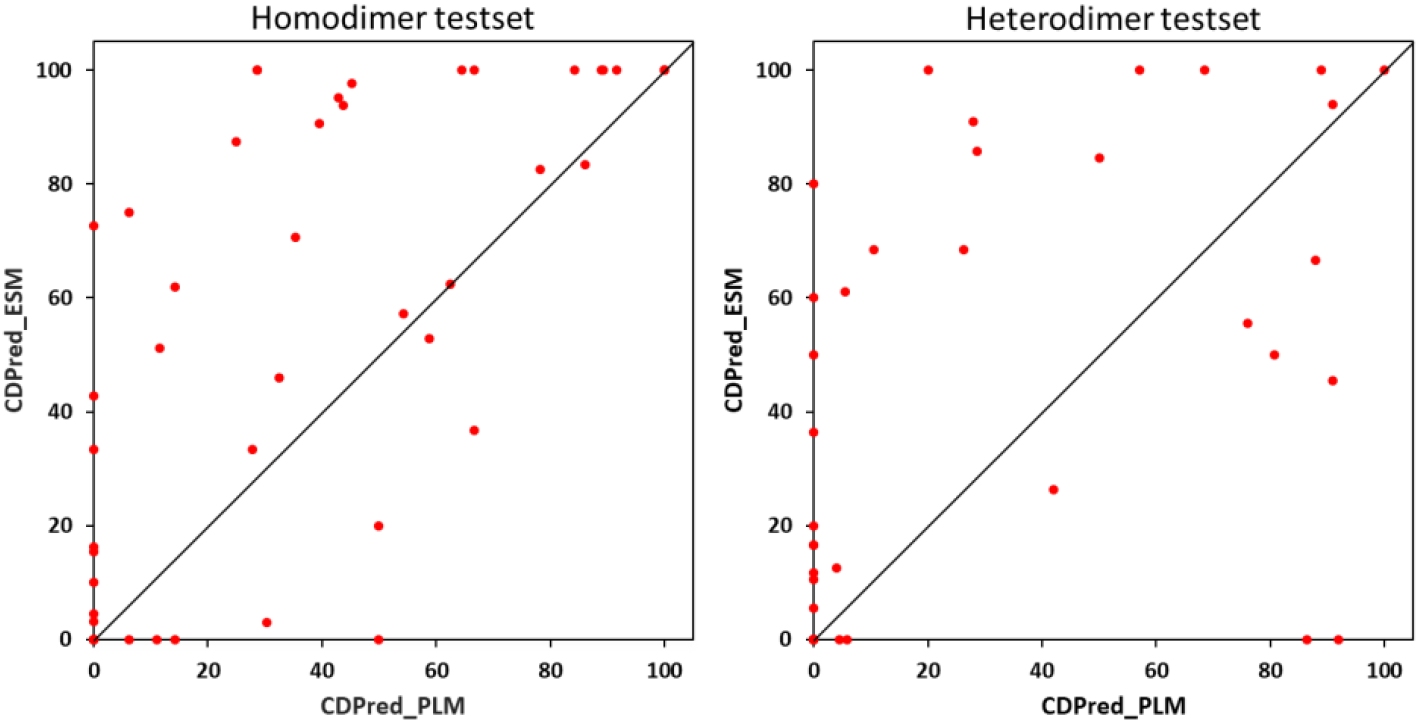
The plot of the top L/10 contact prediction precision (%) of CDPred_ESM against CDPred_PLM for each dimer in the homodimer and heterodimer datasets, respectively. CDPred_ESM has higher precision (dots above the diagonal) for most dimers.

### 2.4 Impact of intra-chain distance information on inter-chain distance prediction

Intra-chain residue-residue distance information of monomers is often used as an input for inter-chain distance/contact prediction. But it is unclear what kind of intra-chain distances may be useful for inter-chain distance/contact prediction. To study this issue, we prepare three kinds of intra-chain distance maps as the input features for CDPred to predict the inter-chain distance map. The first is the full ground-truth intra-chain distance map extracted from the true tertiary structure of monomers (denoted as **FullMap**). The second one is the ground truth distance map that excludes those distances less than 8Å (denoted as **NoContact)**, which can be considered as only keeping non-contact intra-chain distance information (large distances). And the last one is the ground true distance map that excludes those distances greater than 8Å (denoted as **OnlyContact**), which can be considered as only keeping contact intra-chain distance information (small distances). It is worth noting that contact information (small intra-chain distances between residues) is much more important than non-contact information (large intra-chain distances between residues) for determining the tertiary structures of monomers.

The precision of the top L/2 contact predictions using the three kinds of intra-chain distance maps above with CDPred on the two homodimer test datasets and two heterodimer datasets are shown in **Figure 4**. It is interesting that the NoContact intra-chain distance map is much more useful for inter-chain distance map prediction than the OnlyContact intra-chain distance map, indicating that large distances between residues in intra-chain distance maps are much more important for inter-chain contact prediction than small intra-chain distances between residues. The NoContact intra-chain distance maps work even slightly better for inter-chain distance prediction for homodimers than the FullMap intra-chain distance maps, but they perform worse for heterodimers. The same phenomenon is observed when the predicted monomer distance was fed into CDPred for inter-chain distance prediction (data not shown). In summary, the intra-chain contact information is much less informative for the inter-chain distance prediction, but the intra-chain non-contact information plays a critical role in the inter-chain distance prediction. This may be partly due to that most inter-chain contacts involve residues on the surface of monomers that do not have many intra-chain contacts. For homodimers, removing the intra-chain contact information from the input slightly improves the accuracy of the inter-chain distance prediction may be due to that reducing the largely irrelevant part of the input enhances the prediction capability of the deep learning method.

**Fig. 4.**
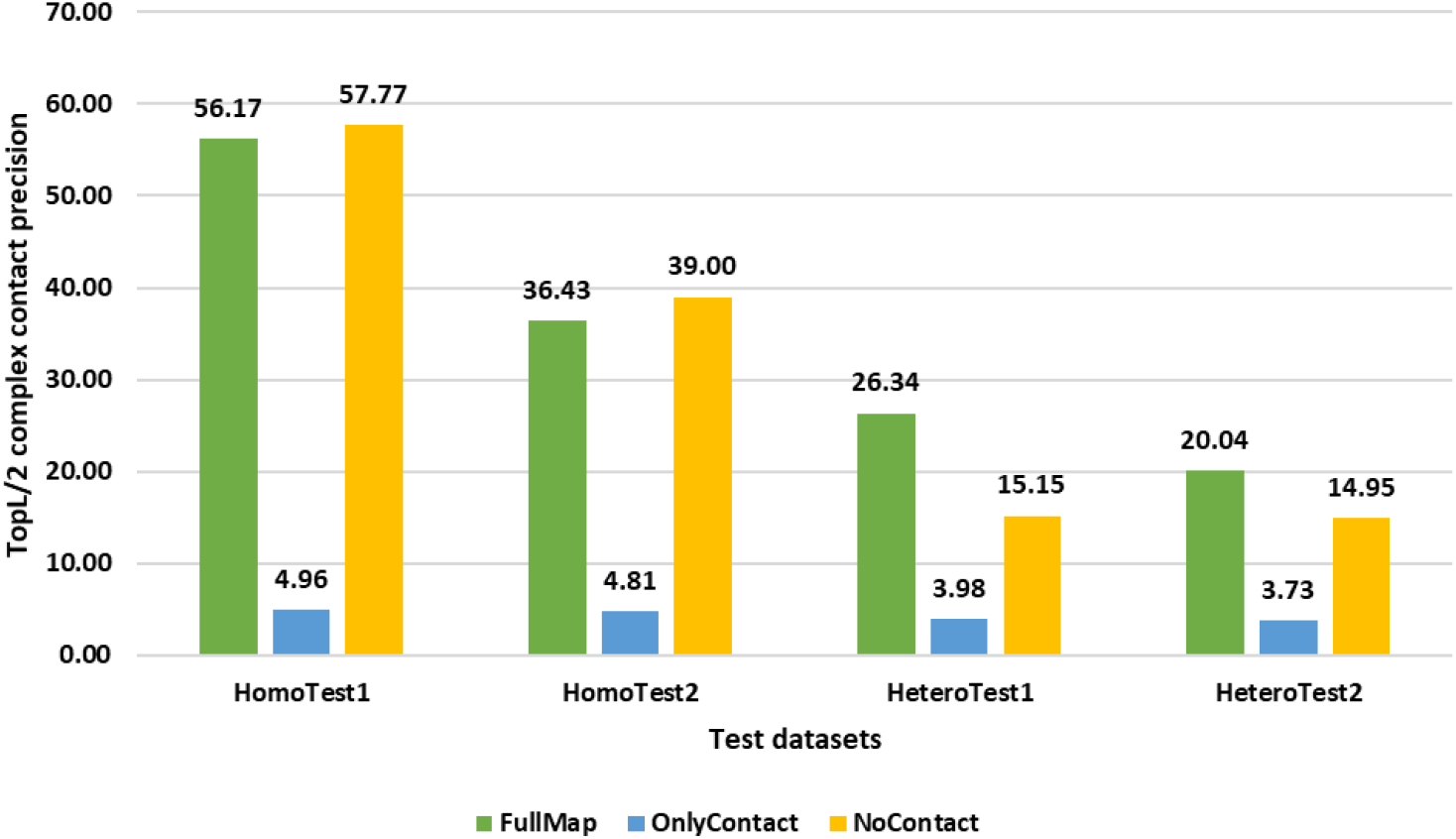
The comparison of the inter-chain contact prediction results with the FullMap intra-chain distance map, the OnlyContact intra-chain distance map, and the NoContact intra-chain distance map as input in terms of top L/2 contact prediction precision (%) on the four test datasets.

### 2.5 High correlation between the precision of inter-chain contact predictions and predicted probability scores

The previous work on the intra-chain distance prediction (Guo, et al., 2021) shows that the intra-chain distance prediction accuracy and predicted probability scores have a strong correlation, which can be used to select predicted intra-chain distance maps. Here, we investigate if the similar correlation exists in the inter-chain distance prediction. **Figure 5** is a plot of the precision of top L/5 inter-chain contact predictions and the average of their probability scores for each target in the four test datasets. The correlation between the top L/5 inter-chain contact precision and the average predicted probability score is 0.7345. The high correlation suggests that the probability of inter-chain contacts predicted by CDPred can be used to estimate the confidence of the inter-chain prediction.

**Fig. 5.**
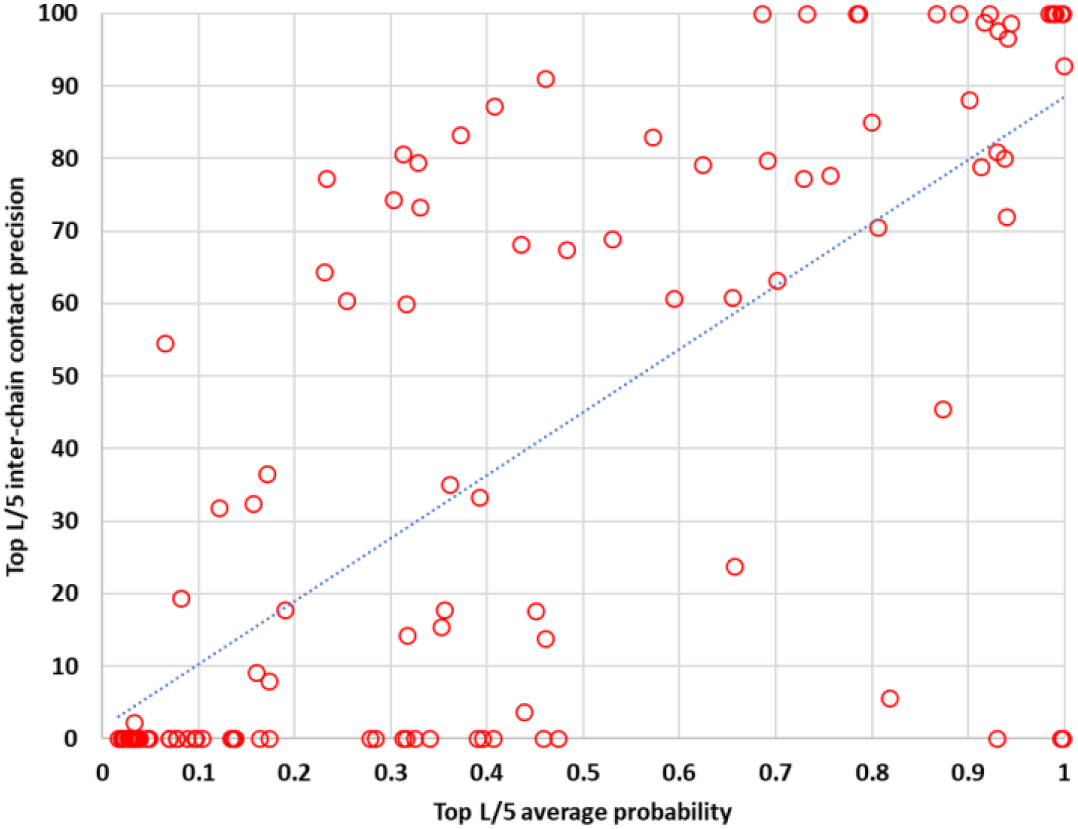
The plot of the precision of top L/5 inter-chain contact predictions against their average contact probability predicted by CDPred for each target in the four test datasets (HomoTest1, HomoTest2, HeteroTest1, HeteroTest2).

### 2.6 An interesting inter-chain distance prediction example

Typically, when the MSA is shallow, the precision of inter-chain distance prediction is low due to the lack of information. However, CDPred still can accurately predict inter-chain distance for some targets with shallow MSAs. **Figure 6** shows such a CASP13 homodimer target T0991. Its MSA has only one sequence. The TM-score (Zhang and Skolnick, 2004) of the tertiary structure of the monomer of T10991 predicted by AlphaFold2 is 0.3104, indicating the predicted tertiary structure fold is not correct. However, the precision of top L/10, top L/5, and top L/2 inter-chain contacts derived from the distance map predicted by CDPred is 72.73%, 68.18%, and 56.36%, respectively, which is high. **Figure 6a** and **6b** show the intra-chain distance maps of the AlphaFold predicted tertiary structure and the true tertiary structure of the monomer, **Figure 6c** shows the inter-chain contact map predicted by CDPred, and **Figure 6d** the true inter-chain contact map. The predicted inter-chain contact map accurately recalls a large portion of the true inter-chain contacts.

**Fig. 6.**
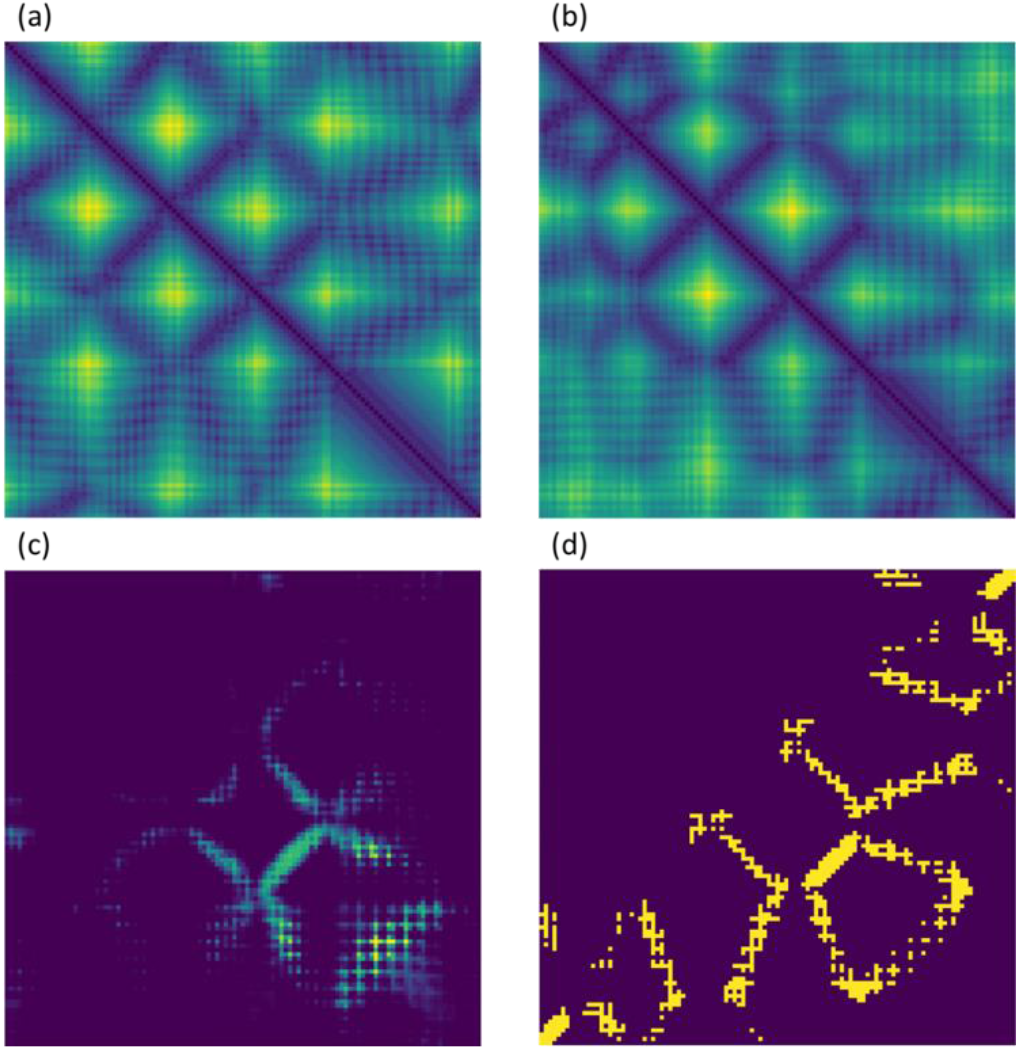
The prediction for homodimer T0991 with a shallow MSA. **(a)** The intra-chain distance map of the monomer is predicted by AlphaFold. **(b)** The true intra-chain distance map of the monomer. **(c)** The inter-chain contact map predicted by CDPred. **(d)** the true inter-chain contact map.

## 3. Conclusion

We developed a new 2D attentive neural network method (CDPred) to predict inter-chain distance maps for both homodimers and heterodimers using multiple sequence alignments and predicted tertiary structures of monomers as input. CDPred substantially outperforms existing methods. We also show that the co-evolutionary features extracted from MSAs by MSA transformer are more effective than those extracted by traditional statistical optimization tools. We discover that the non-contact intra-chain distances (large intra-chain distances) of monomers are much more useful for the inter-chain distance prediction than the contact intra-chain distance (small intra-chain distances). The inter-chain contact prediction accuracy is generally much higher for homodimers than for heterodimers, largely due to the higher depth of the MSAs for the former than for the latter. Therefore, improving the quality of the MSAs for heterodimers is important for advancing the inter-chain distance prediction.

## 4. Materials and Methods

### 4.1. Attention-based neural network architecture

Figure 6 illustrates the overall architecture of CDPred based on the channel-wise and spatial-wise attention mechanisms. CDPred takes the tertiary structures of monomers of a dimer as input and extracts the monomer sequences and intra-chain distance maps. For homodimers, since the sequences of the two monomers of a homodimer are the same, only one monomer tertiary structure is used as input. The monomer sequences are used to search the protein sequence databases to generate MSAs of dimers, which are used to generate residue-residue co-evolutionary scores, row attention maps, and position-specific scoring matrix (PSSM) as input features (see Features Subsection 4.2 for details). The complete input for CDPred is the concatenation of all the input features.

**Fig. 6.**
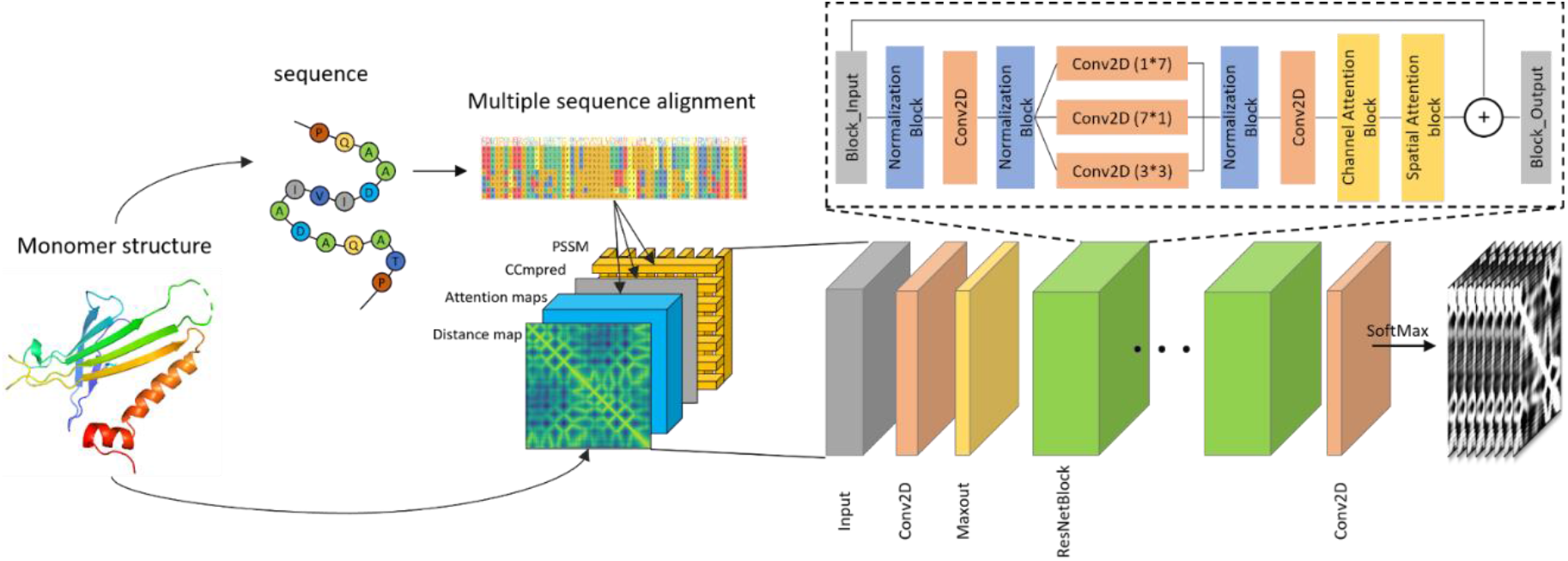
Overview of the CDPred architecture. CDPred simultaneously uses the tertiary structural information (i.e., intra-chain distance map of monomers), sequential information (PSSM), and residue-residue co-evolutionary information (i.e., co-evolutionary scores calculated by CCMpred and attention maps by MSA transformer) as input to predict inter-chain distance maps. The dimension of the input for homomer dimer is L*L*186 (L is the length of the monomer sequence), while the dimension of the input for heterodimer is (L1 + L2) x (L1 + L2) * 186 (L1 and L2 are the length of the two different monomers in the heterodimer). Each of the two output matrices has the same dimension as the input except for the number of output channels. The number of the output channels of the output layer is 42, storing the predicted probability of the distance in 42 distance bins. Two output matrices are generated, representing the two kinds of predicted inter-chain distance maps.

The input features stored in 2D tensors of multiple channels are first transformed by a 2D convolutional layer, followed by a Maxout layer (Goodfellow, et al., 2013) to reduce the dimensionality. The output of the Maxout layer is used as input for a series of deep residual network blocks empowered by the attention mechanism. The residual network has been widely used in computer vision and protein intra-chain distance and contact prediction(Li, et al., 2019; Wang, et al., 2017; Wu, et al., 2020). Here, we combine the residual connection with other useful components to construct a residual block, which includes the normalization block (called RCIN) consisting of row normalization layer (RN), column normalization layer (IN) (Mao, et al., 2019), and instance normalization (IN) (Ulyanov, et al., 2016) for normalizing the feature maps, a channel attention squeeze-and-excitation (SE) block (Hu, et al., 2018) for capturing the important information of different feature channels, and a spatial attention block (Woo, et al., 2018) that captures signals between residues right after the channel attention block. Following the residual blocks, a 2D convolutional layer with the softmax function is used to classify the distance between any two residues from two monomers in a dimer into 42 distance bins (i.e., 40 bins from 2 to 22 Å with a bin size of 0.5 Å, plus a 0-2 Å bin and a >22 Å bin). Two kinds of inter-chain residue-residue distance are predicted at the same time: (1) the distance between the two closest heavy atoms from two residues used by most existing works in the field and (2) the C_b_-C_b_ distance between two residues used by some recent works (e.g., GLINTER), resulting in two kinds of distance maps predicted.

### 4.2. Features

The input features of CDPred contain (1) the tertiary structure information of monomers in the form of intra-chain distance map, (2) pair-wise co-evolutionary features, and (3) sequential amino acid conservation features, which are stored in a L x L x N tensor (L is the length of the sequence of a monomer for a homodimer or the sum of the length of two monomers (L1 + L2) for a heterodimer). N is the number of feature channels for each pair of residues.

#### Tertiary structure information of monomers

The protein tertiary structure information of a monomer in a dimer is represented as an intra-chain distance map storing the distance between C_b_ atoms of two residues in the monomer. For a homodimer, an intra-chain distance map (L x L x 1) computed from the tertiary structure of only one monomer is used. For a heterodimer, two intra-chain distance maps (L1 × L1 × 1 and L2 x L2 × 1) of the two monomers in the heterodimer are computed from their tertiary structures and added as the top left submatrix and the bottom right submatrix of the input distance map of the dimer of (L1 + L2) x (L1 + L2) × 1 dimension. The values of the other area of the input distance map of the heterodimer are set to 0. In the training phase, the true tertiary structures of monomers in the dimers are used to compute the intra-chain distance maps above. During the test/prediction phase, the tertiary structures of monomers predicted by AlphaFold are used to generate the intra-chain distance maps as input. Using predicted tertiary structures as input is more challenging but can more objectively evaluate the performance of inter-chain distance prediction because in most situations the true tertiary structures of the monomers are not known. A predicted tertiary structure also corresponds to an unbound tertiary structure, a term commonly used in the protein docking field.

#### Co-evolutionary features

MSAs are generated for homodimers or heterodimers as input for the calculation of their co-evolutionary features. To challenge the deep learning method to effectively predict inter-chain distance maps from noisy inputs, in the training phase, we use the less sensitive tools or smaller sequence databases to generate MSAs, but in the test phase, we use state-of-the-art tools and larger databases to generate the requisite MSAs. Specifically, in the training phase, for a homodimer, we use PSI-BLAST (Bhagwat and Aravind, 2007) to search the sequence of a monomer against Uniref90 (2018-04) (Mirdita, et al., 2017) to generate the MSAs, and for a heterodimer, we follow the procedure in FoldDock (Bryant, et al., 2022) using the HHblits (Remmert, et al., 2012) to search against Uniclust30 (2017-10) to generate the MSA for each of the two monomers and then pair the two MSAs to produce an MSA for the heterodimer according to the organism taxonomy ID of the sequences.

In the test stage, for a homodimer, we use HHblits to search the sequence of a monomer against the Big Fantastic Database (BFD) (Steinegger, et al., 2019) and Uniclust30 (2017-10) respectively to generate two MSAs for a single chain of the homodimers, which are used to generate input features separately to make two predictions that are averaged as the final predicted distance map; for a heterodimer, an MSA is generated by the same procedure used in EvComplex2 (Green, et al., 2021), which applies the jackhammer to search against Uniref90 (2018-04) to generate one MSA for each of the two monomers and then pairs the sequences from the two MSAs to produce an MSA for the heterodimer according to the highest sequence identity with the monomer sequences in each species. The MSA for a homodimer or a heterodimer is used by a statistical optimization tool CCMpred (Seemayer, et al., 2014) to generate a residue-residue co-evolutionary score matrix (L x L x 1) as features and by a deep learning tool MSA transformer (Rao, et al., 2021) to generate residue-residue relationship (attention) matrices (L x L x 144) as features. L is the number of the columns in MSA.

#### Sequential features

The sequence profile (i.e., position-specific scoring matrix (PSSM)) of protein generated by the PSI-BLAST search above contains the residue conservation information. The PSSM of a monomer in a homodimer or the vertical concatenation of two PSSMs of two monomers in a heterodimer in the shape of L x 20 is tiled (i.e., cross concatenated element by element) to generate sequential features of dimensionality L x L x 40.

#### 4.3. Training procedure

The deep neural network uses the input features above to predict a heavy atom distance map and a C_b_ distance map of shape L x L x 42. The 42 channels store the probability of a distance between two residues in 42 distance bins. The predicted inter-chain distance maps are compared with their true counterparts to calculate the cross-entropy loss to adjust the weights during training. For a heterodimer (L = L1 + L2), an output distance map of dimension (L1 + L2) x (L1 + L2) x 42 contain both inter-chain distance predictions and intra-chain distance predictions. Only inter-chain distance predictions are used to calculate the cross-entropy loss to train the networks, while the intra-chain distance predictions are ignored.

#### 4.4. Datasets and evaluation metrics

We use the DeepHomo training dataset (Yan and Huang, 2021) to train the homodimer inter-chain distance predictor. The whole dataset includes 4,132 homodimeric proteins with C2 symmetry. And after removing proteins that have >=30% sequence identity with the blind test datasets (HomoTest1 and HomoTest2) consisting of the targets of the CASP/CAPRI experiments, 4129 homodimers are left as training, validation, and internal test data. The same as DeepHomo, we select 300 of them as the validation data and 300 as the internal test data and use the rest as the training data. The test dataset used by DeepHomo that contains 28 targets collected from the CASP10-13 experiments is used as one blind homodimer test dataset (**HomoTest1)**. Another test dataset used by GLINTER (Xie and Xu, 2022), which includes 23 homodimer targets collected from the CASP13 and 14, is used as the other blind homodimer test dataset (**HomoTest2**). The two blind test datasets have six common targets.

For heterodimers, we use the heterodimers in Apoc (Gao and Skolnick, 2013) to create the training, validation, and internal test datasets. After filtering out similar sequences at the 40% sequence identity threshold and removing the sequences with >=30% sequence identity with the blind test datasets (HeteroTest1 and HeteroTest2), 3955 heterodimers are left. We randomly select 3576 of them as the training data, 198 as the validation data, and 181 as the internal test data. The test dataset used by GLINTER that contains nine heterodimer targets from the CASP13 and CASP14 experiments in conjunction with the CAPRI experiments is used a blind test dataset (**HeteroTest1**). To create a larger blind test dataset, we collect the heterodimer released between 09-2021 and 11-2021 in the PDB. After filtering out similar sequences at a 40% sequence identity threshold and excluding sequences with >1000 residues targets, 55 heterodimers are left to create another blind test dataset (**HeteroTest2**).

Since the external methods GLINTER and DeepHomo predict inter-chain contacts instead of inter-chain distances, to fairly compare CDPred with them, we use the precision of contact prediction as the evaluation metric. Specifically, the precision of top 5, 10, L/10, L/5, L/2, and L contact predictions (L: length of a monomer in homodimers or length of the shorter monomer in heterodimers) is computed and compared. The similar metric is also widely used in evaluating intra-chain contact prediction. Because DeepHomo and GLINTER predict inter-chain contacts at an 8Å threshold, we use the same threshold to convert the distance maps predicted by CDPred into the binary contact map. A predicted inter-chain contact is correct if the minimal distance between the heavy atoms of the two residues is less than 8Å.

## Acknowledgements

The work was partly supported by the Department of Energy, USA (grant nos.: DE-AR0001213, DE-SC0020400 and DE-SC0021303), National Science Foundation (grant no: DBI1759934 and IIS1763246), and National Institutes of Health (grant no.: R01GM093123 and R35GM118039).

## Code availability

The code of CDPred is available at : https://github.com/BioinfoMachineLearning/CDPred

## Notes

https://github.com/BioinfoMachineLearning/CDPred

https://zenodo.org/record/6647564#.Yq-qi-zMKUn

### Competing Interest Statement

The authors have declared no competing interest.

